# Biological insights and methodological challenges learned from working with a diverse heterotrophic marine bacterial library

**DOI:** 10.1101/2025.11.25.689180

**Authors:** Waseem Bashir Valiya Kalladi, Melisa Osborne, Helen Scott, Franziska Kratzl, Carolina Alejandra Martinez Gutierrez, Elena Forchielli, Natasha Gurevich, Joseline Velasquez-Reyes, Krupa Sampat, Luca Zoccarato, Konrad Herbst, Osnat Weissberg, Hans-Peter Grossart, Daniel Segrè, Daniel Sher

## Abstract

**Background:** Organized collections of bacterial strains can help bridge the gap between studying model organisms and communities, through comparative experiments between genetically and phenotypically diverse strains. We describe the establishment and initial characterization of a library of 62 marine heterotrophic bacteria, selected to represent a significant fraction of the genome-encoded functional diversity and a wide range of known phytoplankton-bacteria interactions. We focus on important but often undiscussed aspects of collecting and maintaining such a library, verifying strain identity, and applying classical microbiological methods across diverse strains.

**Results:** Cultured strains contain up to hundreds of mutations compared with the reference genomes, with non-synonymous mutations in rpoB and/or rpoC genes observed in ∼15% of the cultures. Most strains grow well at 25°C, but the dependence of growth rate on temperature and the width of the temperature niche vary between strains in a systematic manner. We describe steps towards designing a universal, defined, minimal media for marine bacteria, revealing that growth inhibition on amino acids and peptides by carbohydrates is widespread. Cell counts obtained from flow cytometry and colony plating differ systematically, as do different methods to assess motility. Finally, we discuss traits potentially related to microbial interactions such as hemolysis, biofilm formation, and antibiotic resistance. Gammaproteobacteria such as *Alteromonas*, *Pseudoalteromonas,* and *Vibrio* reveal consistently robust growth, and activity, perhaps explaining why these clades are well-explored.

**Conclusion:** Explicitly discussing the insights and challenges of working with strain libraries will pave the way to robust, reproducible, and generalizable mapping of bacterial traits across diversity.

## INTRODUCTION

Bacterial diversity is immense. Environments such as the oceans, soil and the human gut are estimated to contain between 10^6^ and 10^10^ distinct bacterial species, each with its own set of genome-encoded traits ^1–3^. Culture-independent methods such as metagenomics have begun to uncover this diversity by identifying which genes are found in which organisms and how the presence of these organisms and genes varies in time and space ^4–6^. However, in order to understand how genes encode phenotypes, studies with laboratory cultures are needed^7–9^. There are tens of thousands of bacterial strains in culture worldwide, maintained by individual research groups or culture collections, which have provided the foundations for our understanding of bacterial physiology and ecology. Comparing these strains, through the use of bacterial strain libraries, can then allow us to understand how bacteria differ from each other^7,10–12^.

Bacterial strain libraries are typically designed to represent culturable diversity within an ecosystem or to address a specific research question^10,13–16^. Indeed, large and diverse culture collections have enabled scientists to characterize and classify the nature of bacterial interactions to understand the factors governing the classification^7,10,11,17^, understand the metabolic interaction dynamics within a eukaryotic host community^14,18,19^, search for metabolites such as antibacterials or surfactants^20,21^, or understand metabolic requirements and preferences^13,15^.

Isolating, maintaining, and using diverse bacterial culture collections comes with challenges. The handling of strain culture libraries is time and labor-intensive, and the potential exists for operational mistakes such as (cross-) contamination while working with them^22^. Additionally, long term maintenance of laboratory strains might result in the accumulation of mutations^23,24^, resulting in changes in cell phenotype ^25–27^. Yet, to date, there are few studies that discuss, in a detailed and systematic manner, the process of collecting and maintaining bacterial libraries. Additionally, due to the high throughput nature of working with dozens or hundreds of isolates, and the role of statistics in identifying patterns, some of the inherent phenotypic variability is masked when data are presented. These “outliers” can be challenging to deal with yet can represent real facets of the richness of microbial life.

Here, we present insights from the process of assembling a library of marine heterotrophic bacteria, which was designed primarily to study how diverse marine bacteria interact with phytoplankton. We discuss the process of selecting the strains to include in the library, based on their genomic content, collecting them, verifying their identity and propagating them. We describe the growth of these strains at different temperatures and with different media, allowing us to reflect on the variability of cell growth and replication in relation to phylogeny. We further compare different methods for cell counting, highlighting consistent differences between these methods that challenge inter-strain comparisons. Finally, we present a preliminary characterization of several phenotypes related to the interactions between bacteria and other organisms (hemolysis, antibiotic resistance, motility and biofilm formation), using relatively simple and widely used tests. A common thread is that classical microbiological methods can be applied across diverse strains, yet there are challenges involved. Some of these challenges can be partly explained or potentially addressed by revisiting the basic principles of each method. In addition to providing a valuable starting point for future in depth characterization of the various aspects of marine heterotrophs (e.g., their diversity and their ecological interactions), our analysis provides a set of guidelines which may help initiate and expand future efforts for standardized phenotyping of project- or environment-specific bacterial strain libraries.

## RESULTS AND DISCUSSION

### Selecting the strains and assembling the library

For a strain library to be useful, it needs to fulfill several, potentially contrasting, roles. Firstly, it needs to represent, at least to some extent, the group of organisms or environment being studied, yet environments can vary over time and space, containing thousands of organisms. At the same time, the library must be tractable. If the library is designed for use by labs without specialized equipment (e.g. robotic handlers) it needs to be relatively small (several hundreds of organisms at most^11,15^), and the organisms need to be relatively easy to work with and grow under a set of standardized conditions. Thus, any strain library is inherently a compromise. Secondly, in order to infer a potential relationship between genotype and phenotype, the strains in the library need to have high quality sequenced genomes that were annotated in a standard manner. Finally, it will be helpful if the library contains well-studied organisms, as this will help relate comparative studies across the diverse strains and in-depth studies of specific models.

With these concepts in mind, we assembled a strain library ultimately comprising 62 heterotrophic marine bacteria based on a comparative genomic study and on previous studies of phytoplankton-bacteria interactions (Figure 1A, Supplementary Figure 1, Supplementary Excel File ^28^). We previously analyzed the genomes of 473 marine bacteria and clustered them into “Genome Functional Clusters” (GFCs), each of which was predicted to have similar metabolic potential (KEGG modules) and similar capacities to interact with other organisms. Through an exhaustive search of published literature, as well as the catalogues of culture collections such as the German collection of Microorganisms and Cell Cultures (DMSZ) and the American Type Culture Collection (ATCC), we identified strains representing as many of the GFCs as possible (see below). To this list we added ∼20 strains selected based on previous studies of their interactions with phytoplankton^11^, as well as two well-studied reference organisms (two *E. coli* strains, MRE600 and ATCC25922).

**Figure 1:**
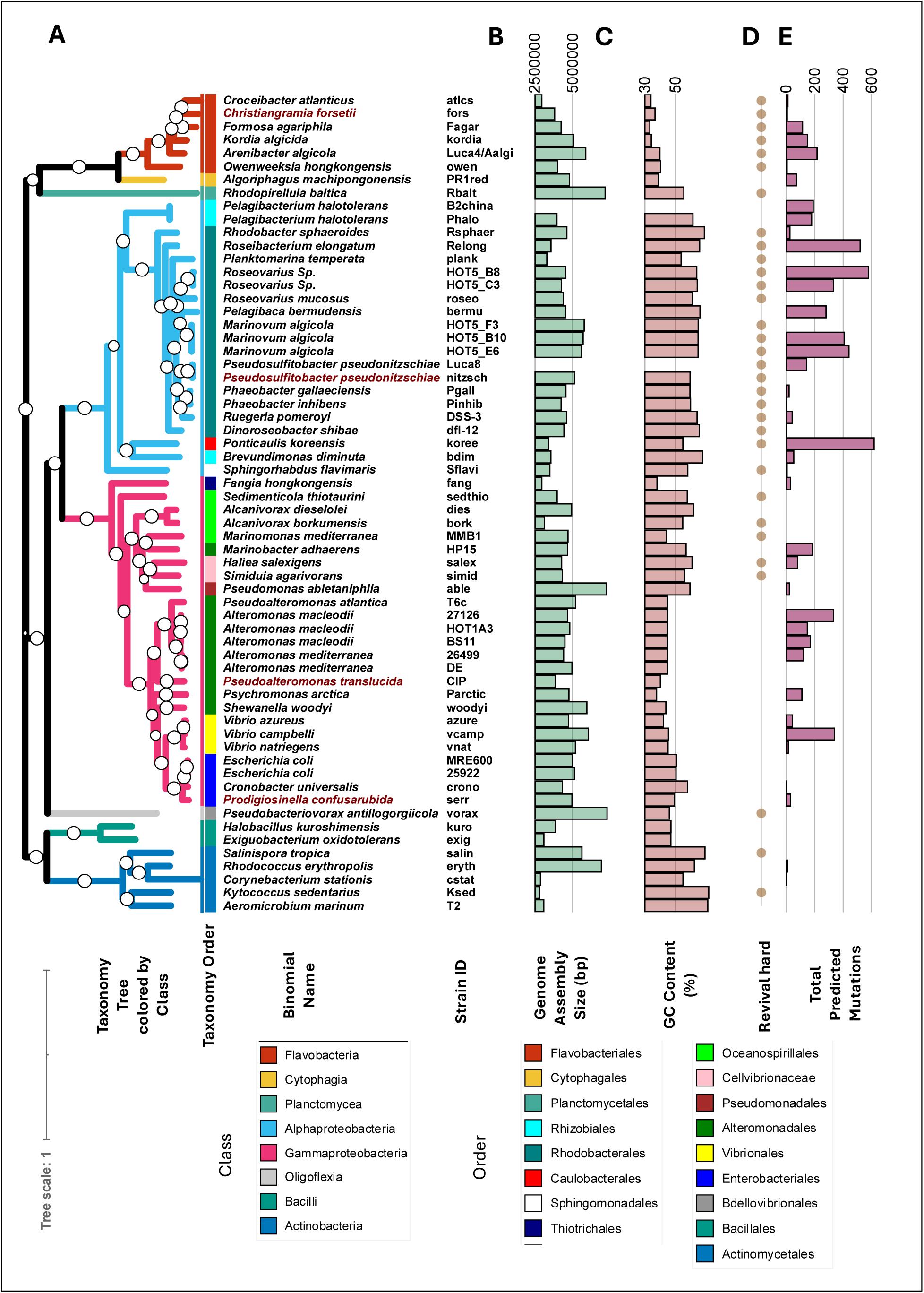
The library structure. A) Phylogenomic tree of the strains under study colored by taxonomic class. Bootstrap values are shown in white circles, color strip shows taxonomic order. The binomial names of strains that were renamed during the period of this study are highlighted in brown. B) Genome size in base pairs. C) Genome GC content. D) Strains that were hard to revive at least once. These strains required using large volumes of the glycerol stock to revive a handful of colonies. E) Number of observed mutations per genome, including both SNPs and indels.

We chose to limit ourselves to strains classified as Bio Safely Level 1, and thus the library does not contain pathogens such as *Aeromonas*, *Clostridium,* and some *Vibrio* that can be found in marine environments. We also required the strains to grow on Marine Broth (MB) or Marine Agar (MA), and at room temperature, meaning that the library does not contain abundant clades such as *Pelagibacter* (SAR11). Many of the strains in this initial list were no longer available, due to technical issues (e.g. “Our -80 freezer crashed”) or lab closure, underscoring the importance of depositing well-studied strains in professional culture collections. 69 strains were ultimately received, of which 4 were removed due to discrepancies between the expected strain and the results of 16S or genome sequencing (see below), and another 3 revealed very slow or irreproducible growth on MB or MA. Of the final 62 strains, approximately three quarters were obtained from culture collections, and the rest from seven individual labs (Supplementary Figure 1A). They represent approximately half of the GFCs ^28^ (Supplementary Excel File).

The range of genome sizes and GC content of the collected strains (traits which were not taken into account when we selected the strains) was slightly smaller than those of the 473 genomes in the initial analysis (2.7-7.2 vs 1.2-9.8 Mbp and 33.3-71.6% vs 27.9-71.9% GC). This is likely due to our choice not to include fastidious organisms with highly streamlined, often GC-poor genomes such as SAR11. There was no correlation between genome size and phylogeny, whereas the GC content tended to be higher in Alphaproteobacteria and Flavobacteria (Figure 1B, C). Most of the strains in the library were originally isolated from seawater, with the rest obtained from sediment, microalgal cultures, macro-organisms and other environments (Supplementary Figure 1B). Most were isolated from coastal locations, and from surface waters. However, for most of the strains, the exact location and date where they were isolated are unknown (64% and 58%, respectively, Supplementary Figure 1B). This will complicate potential studies attempting to link genotype or phenotype to a specific location or season^29,30^, underscoring the importance of collecting and reporting as much metadata as possible when describing a new isolate.

### Strain preservation and validation

When the 69 strains arrived in our labs, they were re-streaked for purity and the resulting cultures were frozen at -80°C in aliquots of 50% glycerol in MB as laboratory archives. Due to reports of inconsistencies between plasmid profiles and genomes of strains that were supposed to be identical^31^, we had planned to re-sequence the genomes of all strains, but circumstantial factors such as the covid pandemic resulted in this validation step not being prioritized. However, after several years we observed that some strains did not revive well from the cryo-preserved stocks, primarily those high GC content (>50%, Figure 1D. Often these strains could only be revived when a large volume of the glycerol stocks was plated, resulting in very few colonies, which raised the concern of genetic bottlenecks and/or contamination. We therefore sequenced the 16S rRNA gene amplicon, identifying two cases where the identity of the strain was not the same as the expected one (*Pseudoalteromonas citrea* and *Roseobacter denitrificans*). One additional strain, *Algoriphagus machipongonensis*, exhibited two colony morphologies (red and white), one of which the latter was identified as a contaminant (*Paracoccus yeei*) and removed. Genome re-sequencing (discussed below) identified another two strains which may have been mixed up, based either on a high number of mutations (*Alteromonas mediterranea* DE1) or on low mapping coverage (*Shewanella denitrificans*). These strains were removed, with the final library comprising 62 strains (Figure 1).

Because strain verification through 16S rRNA amplification or genome resequencing was performed only after several years in our labs, we do not know whether the strains were mixed up or contaminated in our hands, or whether they were already contaminated or incorrect when they arrived from their source. We therefore recommend that the identity of any new strain arriving in a lab, whether or not it is part of a library, be verified. Sequencing the 16S rRNA gene amplicon is likely the most straightforward and rapid way to do so, but the sequence of this gene is highly conserved in some clades, and thus it cannot be used to differentiate, for example, between strains of *Vibrio, Escherichia, Alteromonas* or *Pelagibaca*^32–35^. We also note that bacterial strains are regularly re-named or reclassified (e.g. *Pseudosulfitobacter pseudonitzschiae*^36^ and *Pseudoalteromonas haloplanktis*^37^ in our library), requiring careful book-keeping. Finally, we note that the question of how best to maintain diverse strains for extended periods of time is still unresolved. Glycerol archives of some model organisms such as *E. coli* can be maintained for decades at -80°C, but our experience and that of others suggests this is not the case for many other bacteria ^38,39^. Other solutions, such as storage in liquid nitrogen or lyophilization, should also be considered.

### Genomic differences between library strains and reference genomes

To identify to what extent the strains in our library have accumulated mutations compared to the reference genomes, we re-sequenced 48 strains using Illumina technology, analyzing 42 of them for which coverage was high (>20x). The number of mutations (including single nucleotide polymorphisms, insertions and deletions) ranged from 0 (*Pseudoalteromonas atlantica*, named “t6” in our collection) to 618 (*Ponticaulis koreensis,* koree), with a mean of 148 mutations per genome (Figure 1E). There was no correlation between the number of mutations and the sequencing coverage, and there was no significant difference between the number of mutations in strains we obtained from individual researchers compared with those originating in culture collections (203±171 and 118±163, p=0.12, two-tailed t-test). Intriguingly, there were, on average, twice as many mutations in Alphaproteobacterial genomes compared with those from Gammaproteobacteria (215±216 vs 114±112, Figure 1E), although this difference was not significant (p=0.06, two-tailed t-test). There was also a wide range in the value of dN/dS (from 0.13-10) and in the ratio of total SNPs to total indels (from 0 to 29). These data suggest that genomes of bacterial cultures are likely to contain dozens of genetic differences compared to the published genomes of the same strains. As described above, we do not have detailed information on the time the strains were in culture, or on the conditions under which they were maintained. Thus, we cannot identify the causes of these genetic differences, although we suggest that they may depend to some extent on the quality of the initial genome assembly.

Three genes, *rpoB*, *rpoC* and *ssb1*, were mutated more commonly in the library strains compared to other genes (5, 3 and 3 non-synonymous mutations across 42 strains, respectively, compared with 40 genes mutated in 2 strains and 223 in one strain, (Supplementary Excel File). *rpoB* and *rpoC* are two subunits of the RNA polymerase core enzyme, and mutations in these genes are known to confer resistance to a wide variety of stressors including antibiotics, heat and starvation across multiple bacteria ^40^. The *ssb1* gene encodes a single-stranded DNA binding protein involved in DNA replication, recombination and repair. This gene could also be important for resistance to various stress conditions, although this has been debated ^41^. We speculate that these mutations (as well as potentially other ones) may be induced by the common practice of maintaining bacteria for extended periods as colonies or agar stabs, where they may be exposed to nutrient starvation, cold or other stressors. In line with this, we propose that researchers consider re-sequencing any strain obtained from a collaborator or culture collection, particularly if it is to be used for studies involving response to various stressors.

### Growth rate across experimental temperatures

Growth rate is an important trait in microbial ecology, differing between organisms, and at times determining which strains outcompete others ^42–44^. Multiple approaches are used to infer growth rate in-situ, but studies of growth rate across cultured isolates are critical to separate the effect of different drivers of growth rate (e.g. nutrient concentration, temperature, predation), thus providing context for field studies. Additionally, comparative experiments with multiple isolates could help identify the molecular mechanisms governing growth rates and determine to what extent growth rates affect cell structure ^45^. We measured the growth of the strains in the library on Marine Broth (defined as change in culture turbidity at 600nm (OD_600_), h^-1^) at six temperatures ranging from 10-30°C (Figure 2A). The growth rates varied between 0.0001 (no growth) to 0.27 h^-1^ (0.0024 to 6.48 d^-1^), consistent with rates measured in the field across multiple locations (0.3-10 d^-1^) ^43,44^. Gammaproteobacteria with low GC Content such as the Alteromonadales exhibit both the highest absolute growth rates and the widest thermal niche, growing at rates higher than 0.266 h^-1^ across all temperatures tested (Figure 2A). In contrast, Gammaproteobacteria with higher GC contents, such as Oceanospirillales, as well as most Alphaproteobacteria, tended to have lower growth rates and narrower temperature niches. However, while 25°C was the optimal growth temperature for most strains (Figure 2B), Alphaproteobacteria grew best at 30°C, suggesting we may not have covered their optimum in this experiment.

**Figure 2:**
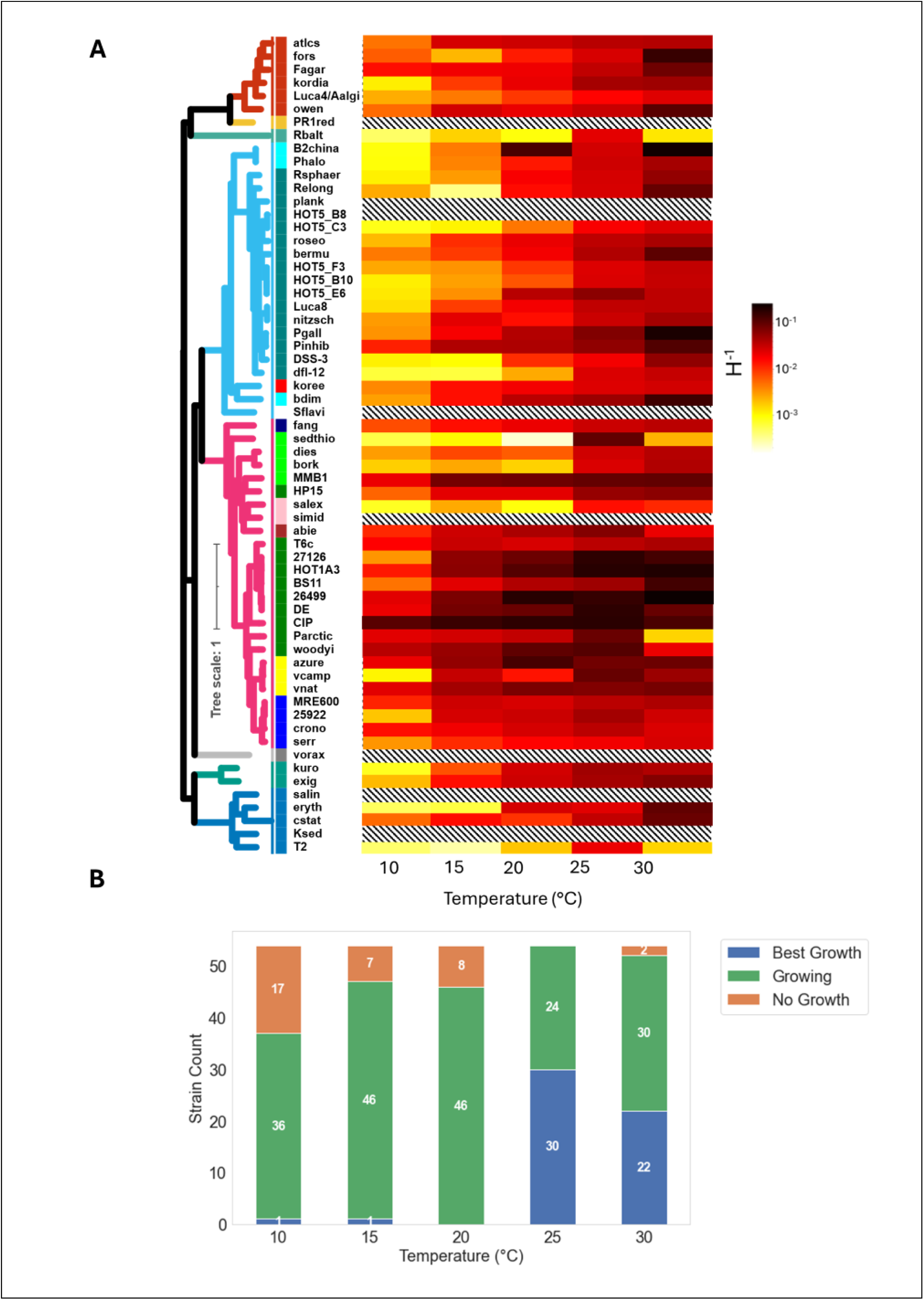
Growth Rates and Temperature. A) Heatmap of the Maximal Growth Rate (h^-1^) calculated for a given strain for each of the tested temperature. White-and-black transverse stripes mark non-tested strains. B) Stacked bar plot of growth at a given temperature classified into non-growing, growing, and best growing. Strains are arranged by same tree in Figure 1.

### Towards a defined, minimal, growth medium

Complex, chemically undefined media such as Luria Bertani Broth (LB, also known as Lysogeny Broth) or Marine Broth are fundamental tools in microbiology, simple to prepare and appropriate for growing a wide range of bacteria. However, because they are not chemically defined, they cannot be used, for example, to determine whether bacteria can grow on specific organic or inorganic molecules as sole sources of energy, carbon, or other elements. Additionally, it is hard to quantitatively assess growth yield or efficiency in such media, as the causes that ultimately limit growth are not always clear (e.g. depletion of specific carbon, nitrogen, phosphorus, or iron sources, depletion of oxygen, or waste accumulation,^46–48^). An ongoing goal of our work with the library is therefore to develop a growth media widely applicable for marine bacteria that has the smallest possible number of chemically defined resources, making it both defined and minimal.

As a first step towards this goal, we previously “re-factored” Marine Broth, replacing peptone and yeast extract as carbon sources with one or a combination of eight molecular classes of organic matter - peptides, amino acids, lipids, disaccharides, organic acids, neutral sugars, amino sugars, and acidic sugars^15^. No single molecular class was able to support the growth of more than ∼65% of the strains, although amino acids and peptides supported the highest amount of biomass growth (maximum OD_600_). We therefore reasoned that combining amino acids or peptides with a defined set of carbohydrates (maltose, trehalose, sucrose, lactate, pyruvate, acetate, citrate, and glycolate) might increase the range of bacteria growing, as well as the yield. We also asked whether whole protein could replace peptides and amino acids, since many bacteria have the ability to degrade extracellular protein^49^. We chose to use a α-casein as a single source for the protein, peptides and amino acids, as it is well-defined and contains all 20 proteinogenic amino acids.

Overall, growth was highest on peptides, followed by amino acids (maximum measured OD_600_, measured every 12h for 4 days, Figure 3A). The slightly higher average growth on peptides rather than amino acids is consistent with our previous results^15^. While some recent studies have begun to systematically identify the substrates of transporters in marine bacteria, including some that take up amino acids^50,51^, much less is known about the presence and specificity of peptide transporters^50,52^. Since peptides are known to be an important resource in some marine communities, better understanding peptide production, uptake and utilization is an important avenue for future research^53^. While the whole protein revealed a lower overall growth yield compared with peptides and amino acids, it still supported significant growth in many strains, suggesting a widespread ability to degrade extracellular protein.

**Figure 3:**
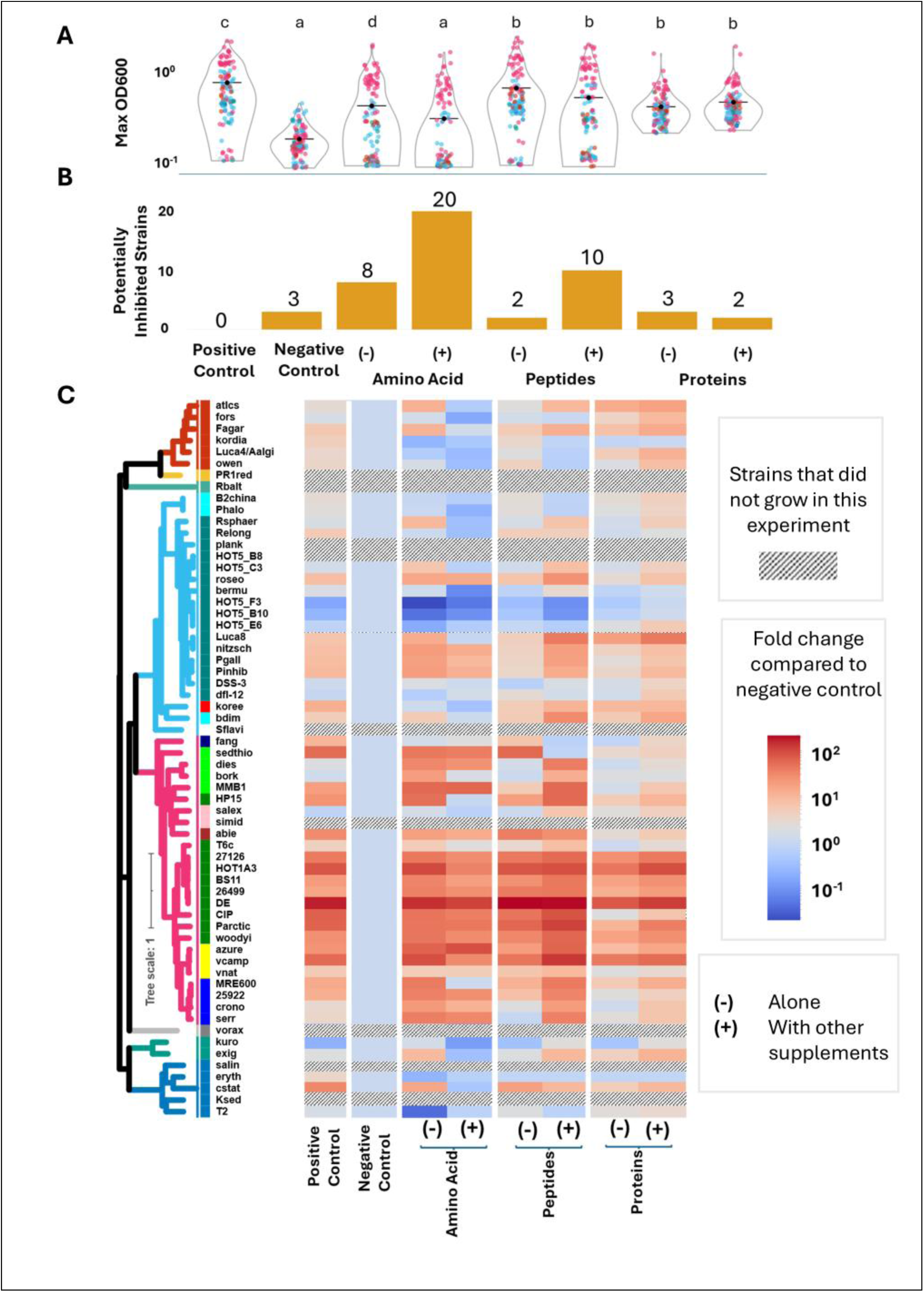
Growth of strains on proteins, peptides and amino acids with and without carbohydrate addition. A) Maximum absorbance at 600nm for each media. Letters above violins represent significance groups from a Kruskal–Wallis test (H = 235.711, p < 0.001) followed by Dunn’s post hoc test with Benjamini–Hochberg false discovery rate correction; media sharing a letter are not significantly different. B) Number of strains that were potentially inhibited (i.e. maximum OD600nm was lower than the blank) on each media. C) Heat map of the log10 fold change in maximal OD600 achieved by each strain on each media. – and + are without and with the addition of carbohydrates.

While the addition of amino acids, peptides or protein enabled growth of some strains, it also potentially inhibited others, i.e. reduced the OD_600_ to that of the blank (not containing any inoculated bacteria), or below. The negative control (media without any carbon source) inhibited only 3/62 strains, a phenomenon observed previously, which suggests many marine bacteria to not die rapidly due to starvation^54^. In contrast, amino acids or peptides inhibited about ∼15% of the strains, suggesting these resources may be toxic to some bacteria when used as sole C sources (Figure 3B). Amino acid toxicity has been studied in several bacteria but usually using model organisms such as *E. coli* and individual amino acids with a much higher concentration rather than a mixture^55,56^.

Unexpectedly, combining resources, i.e. the addition of carbohydrates to peptides or amino acids, actually increased the number of potentially inhibited strains, and non-significantly decreased the growth yield (Figure 3B, C). This effect was not seen when carbohydrates were combined with proteins, which may even have increased the number of strains growing. The addition of carbohydrates seemed to be detrimental primarily to Flavobacteria, Bacilli, Actinobacteria, and most Alphaproteobacteria (the latter with the exception of the genera *Pseudosulfitobacter, Phaeobacter* and *Marinovum*) (Figure 3C). We speculate that the reduction of growth on amino acids and peptides when carbohydrates are added is due to carbon catabolite repression (CCR). First described as “diauxie” in classic papers by Jacques Monod^57^, CCR is a mechanism allowing bacteria to prioritize one resource over another, with the aim of maximizing growth rate^58^. However, we note that CCR is usually described as a (temporary) cessation in growth, rather than inhibition as we observed here. Alternative hypotheses could include osmotic stress from the supplementation^59^, metabolic overflow leading to toxicity by metabolite accumulations^60^, and metabolically wired pH fluctuations^61^.

### Comparing methods of cell counting

The question of how to count bacterial cells in a sample was foundational to quantitative microbiology^62^. Two main approaches have been used for decades: direct counts, e.g. using microscopy or flow cytometry (FCM), and counting colony forming units (CFUs). Each of these methods has its advantages and disadvantages. CFU is often regarded as the benchmark of cell counts, but it is labor intensive, requires a long wait time for colonies to grow and be counted, and not all bacteria produce colonies. FCM is faster than CFU counts (can provide results in minutes), yet is can be difficult to distinguish live cells from dead ones or debris^63–65^. Both methods can be affected by cell aggregation, leading to multiple cells being counted as one. With the advent of molecular techniques other approaches have also been employed, including quantitative PCR (qPCR) using general or species-specific markers^66^, or (in complex communities, and with appropriate calibration), amplicon sequencing^67^. However, molecular methods tend to be time-consuming and expensive, and the results can be affected by differences in DNA copy number, extraction or amplification yields^68,69^. Since cell counts are important in microbiology there have been a few comparisons between methods for specific applications^70–72^, yet there is no standard counting method and, to the best of our knowledge, no direct comparison between methods across diverse bacteria.

Figure 4 shows a direct comparison between counts of bacteria in the library using FCM and CFU (the drop assay, as described in the materials and methods) after 36 hours of growth in Marine Broth. FCM is complicated by a high background staining that partly overlaps the signal of the bacteria themselves and likely originates from a combination of carryover between samples, and particles present in the Marine Broth (Supplementary Figure 2). As a result, no cell counts were available for several cultures even though colonies were observed (e.g. strains 25922, Aalgi and HOT-1A3, see below). For other samples, FCM-based cell counts ranged approximately between 10^9^ and 10^11^ cells/ml. No clear phylogenetic pattern was observed in cell counts (Figure 4B), forward scatter (which is related to cell size) or Sybr Green fluorescence (Supplementary Figure 3, Supplementary Excel File). Indeed, there was no correlation between Sybr Green fluorescence and genome size (or GC content), even though Sybr Green binds nucleic acids. This contrasts with a previous study using a different DNA dye (DAPI) across 4 bacterial isolates^73^.

**Figure 4:**
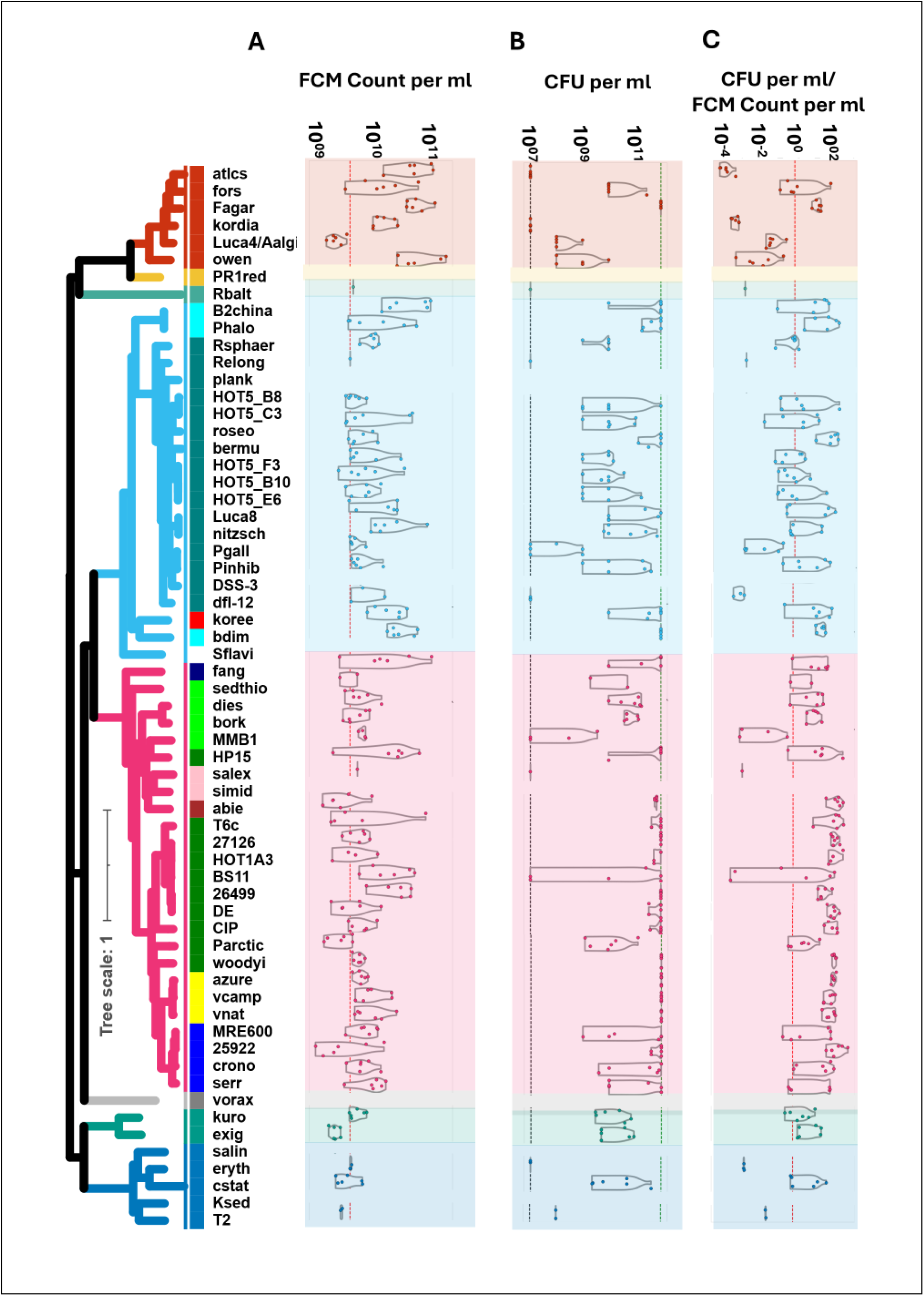
Comparison of cell counts using flow cytometry and colony forming units. A) Flow cytometry counts. B) Colony Forming Units. The upper and lower lines represent cells too numerous to count (Green line) and too low to count (Black line) C) The ratio of CFU per ml/ FCM per ml (B).

The CFU counts revealed a much wider range, between <10^7^ and ∼10^12^ cells/ml (too low and too high to count, respectively). In contrast to the FCM, a clear phylogenetic pattern was observed in the CFU counts, with most Gamma-proteobacteria from the orders Alteromonadales, Vibrionales and Enterobacteriales being too high to count. For many strains, the variation in CFU counts was very high, due to the need to perform 10-fold dilutions in order to enable us to count the entire library without automation (which would require hundreds of plates). The FCM and CFU results were not correlated.

In principle, FCM and CFU counts do not need to agree, since the former counts total cells and the latter only those able to form colonies. If this is correct, the ratio between absolute cell counts and CFUs could be used to infer the fraction of viable cells. For most Alphaproteobacteria this ratio is close to 1, suggesting most cells can form colonies, whereas for 4 out of 6 Flavobacterial strains this ratio is below 0.1, suggesting many cells could be non-viable. However, for many strains in the library the CFU counts were much higher than the FCM ones, sometimes by two orders of magnitude (Figure 4B). The systematic nature of the high CFU/FCM ratio, which was mostly observed in Gammaproteobacteria, suggests that it is due to physiological differences between clades. Underestimation in FCM counts could be due, for example, to systematic differences in the thickness of the peptidoglycan layer or other modifications such as N-glycosylation of Actinomycetales^74^, which could inhibit the binding of the DNA dye used to differentiate cells from debris. In parallel, overestimation of the CFUs could be due to cells (or cell clumps) adhering to the pipette tips used to perform serial dilutions, potentially though systematic differences in, for example, cell surface hydrophobicity. Until the discrepancies between the two methods are resolved, or at least better understood, we suggest that any experiment which needs to be interpreted in light of robust cell counts should employ multiple methods, focusing on results that are not dependent on the method of counting.

### Hemolysis and antibiotic resistance

The ability to produce compounds that can harm other cells, and to resist such compounds, are key traits that could determine how bacteria interact with each other, with phytoplankton cells, and with other hosts. Several previous studies using collections of cultured marine bacteria have consistently shown that the ability to inhibit the growth of other bacteria is quite common, with ∼20-70% of the strains able to inhibit at least one other strain^10,17,34,75^. When inhibition was tested across a broad range of bacteria, Gammaproteobacteria^14^, Alphaproteobacteria or Actinobacteria^75^ tended to be the most inhibitory. Within the Vibrio clade, inhibition was more common between strains that belong to genetically different populations^17^. With this in mind, we assessed two other “interaction” traits that are relatively easy to measure using simple and commonly used methods – hemolysis and antibiotic resistance.

Hemolysis is the ability to produce compounds (hemolysins) that can disrupt the cells of competing organisms or predators, such as protozoa or other microbes, but is usually assessed using human red blood cells in liquid or on plates. Genes encoding putative hemolysin-like toxins are common in the genomes of some marine bacteria, including intracellular symbionts of hydrothermal vent mussels^76^, yet hemolytic activity has typically only been tested in fish pathogens^77^. We used blood agar plates, which are commonly employed in medical microbiology, to categorize the hemolytic activity of each strain into alpha (partial or incomplete hemolysis), beta (complete hemolysis), and gamma (no hemolysis) types (Figure 5A). Complete (beta) hemolysis, which is usually attributed to the production of protein toxins, was observed primarily in Vibrionales, Enterobacteriales, Bacillales and Actinomycetales, orders previously shown to contain hemolytic and often pathogenic organisms^78^. Alpha (incomplete) hemolysis, which results in a greenish/brown discoloration of the media rather than in complete loss of the red color, was observed in most library strains, with no clear phylogenetic pattern, whereas no hemolysis (gamma) was rare. Incomplete hemolysis is not necessarily the result of hemolysin production, since it can also be caused by oxidation of oxy-hemoglobin by hydrogen peroxide^79^. Since hydrogen peroxide and other Reactive Oxygen Species (ROS) such as superoxide are produced by many marine bacteria, this could potentially account for the observed “hemolysis”^80,81^.

**Figure 5:**
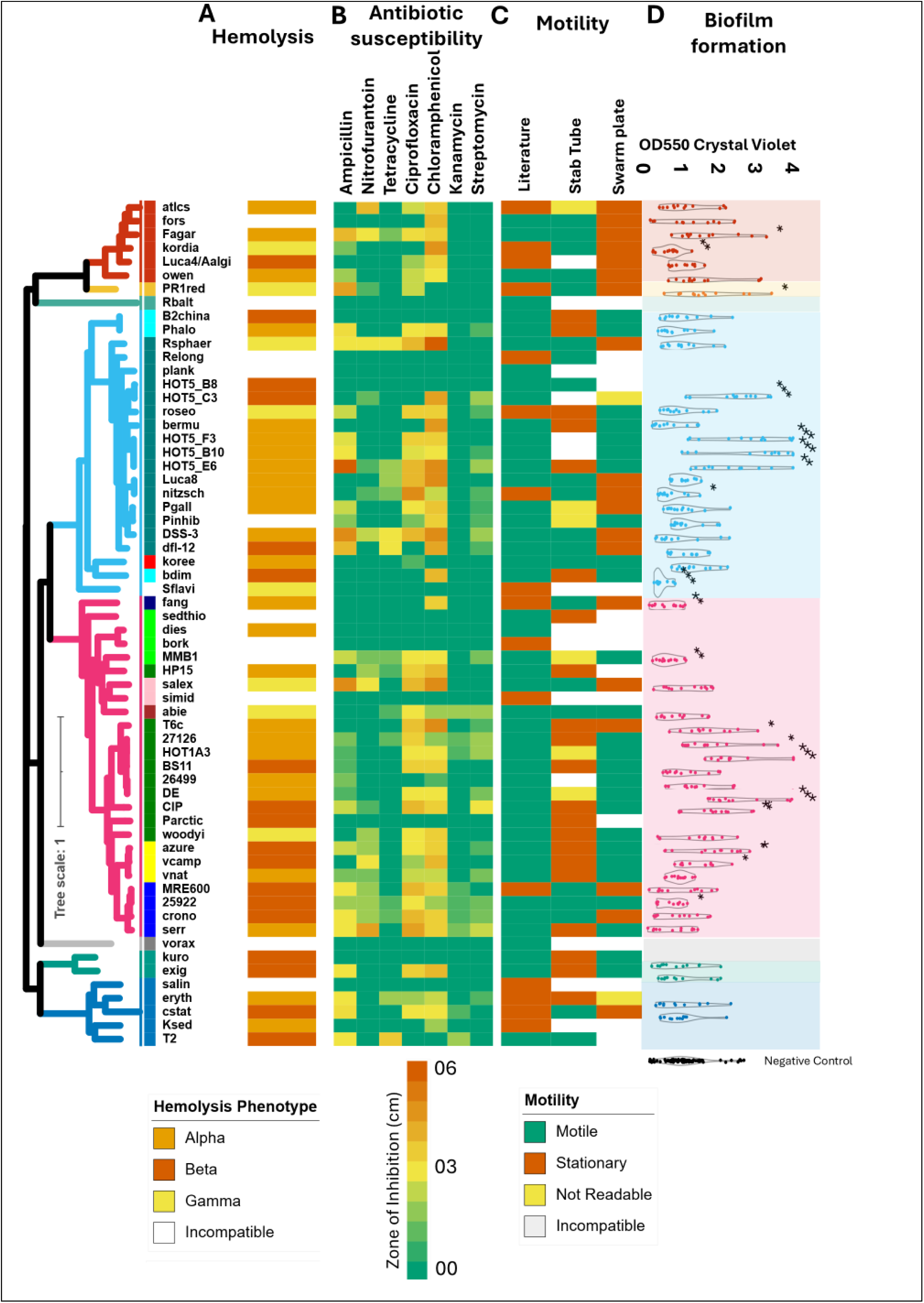
Interactions phenotyping in bacteria. The motility profile of the bacterium shown as described in the literature and assessed by two different methods (A). The hemolysis phenotype on Blood Agar Plates (B), the antibiotic resistance profile accessed by Kirby-Bauer test (C), and the crystal violet test for biofilm formation (D). Statistical significance was assessed using Welch’s t-test (each strain vs. control) with Benjamini–Hochberg correction for multiple comparisons (FDR < 0.05); significance levels are indicated as *p < 0.05, **p < 0.01, ***p < 0.001.

Antibiotic susceptibility was assessed on Marine Agar plates using the Kirby-Bauer assays, which measure a zone of inhibition around a small filter paper containing the antibiotic (Figure 5B). Chloramphenicol, ampicillin, and ciprofloxacin inhibited most strains, consistent with a previous study of arctic bacteria ^78^, with no clear phylogenetic pattern. Strains that were resistant to these antibiotics were dispersed among all clades, suggesting that the resistance mechanisms may derive from horizontal gene transfer^82^. In contrast, most strains were relatively resistant to Tetracycline, Kanamycin and Streptomycin, which all bind to the 30S ribosomal unit, and which are all highly inhibitory to arctic bacteria^78^.

It has previously been shown that marine Gammaproteobacteria are the most resistant clade to inhibition by other bacteria, whereas Bacteroidetes (Flavobacteria and Cytophaga) were most susceptible^10^. We do not observe this pattern in our data, suggesting that either the compounds produced in the study of bacteria-bacteria inhibition are different from those tested here, or that the inhibition phenotype is more complex, involving multiple molecules or multiple mechanisms (e.g. production of antimicrobial compounds together with competition for nutrients).

### Motility and biofilm formation

Many marine bacteria shuttle between living on various particles and swimming freely^83^. The abilities to swim towards, adhere to, and utilize the resources on a particle are thought to be important traits determining the ecological niche of a bacterium, and may have important biogeochemical consequences^83–85^. We therefore asked whether the bacteria in our library are motile and whether they can form adherent biofilms. There are well-established assays for both of these traits, and indeed bacterial motility is often assessed as part of the phenotypic description of a new strain. Yet, as discussed below, these ostensibly simple assays highlight that these two traits are in fact complex and functionally diverse, and that measuring them can be complicated by the phenotypic plasticity of many bacteria.

There are many types of bacterial motility^86^. We chose to classify bacteria into motile and non-motile using two widely accepted methods-the swarm plate test and the stab tube test^87–90^. Both methods rely on measuring the dispersal of bacteria on agar, but they differ in the physical properties of the environment (surface of the agar vs submerged stab into the agar), in the direction of motility (horizontal on plates vs vertical within the stab), and in the oxygen concentrations (aerobic on the plate, gradient of oxygen from aerobic to micro-aerobic or anaerobic in the stab). A literature survey showed that most strains were described as motile, with the exception of most Flavobacteria and Actinobacteria (Figure 5C). However, in >90% of the cases, there were discrepancies between the two motility assays we employed, and therefore also with the literature. Specifically, the Alteromonadales were motile only on plates (consistent with the literature describing them as motile), whereas Flavobacteria were only motile in test stabs, possibly explaining why most were previously described as non-motile (Figure 5C). The three Vibrio species, which were all previously described as motile, were non-motile in the stab assay, despite being facultative anaerobes. Thus, motility in stab cultures may be affected by other environmental factors, besides oxygen concentrations. These results highlight that bacterial motility is strongly dependent on culturing conditions^91,92^.

Once marine bacteria encounter a surface, they often attach to it, forming microcolonies and then biofilms^93^. There are many methods to measure biofilm, each with its own assumptions and caveats^94^. One of the most commonly used assays utilizes crystal violet, which binds to negatively charged molecules in bacterial cells and extracellular matrix components of biofilms^95^. To test the applicability of the crystal violet assay across the diverse library strains, we grew them on marine broth in 96 well plates for 76 hours, observing that some Gammaproteobacteria (primarily Alteromonads) and some Alphaproteobacteria (e.g. Marinovum, and Roseovarius) consistently produced biofilms (figure 5D). However, little biofilm production was observed with Vibrio, or Escherichia, genera that are known biofilm producers^93,96^. One reason for this discrepancy could be the relatively high background signal from the marine broth itself, perhaps due to organic matter precipitating on the plastic surfaces. Perhaps more importantly, biofilm growth is a complex process, which is sensitive to many environmental conditions, including the initial cell density (which we did not measure here), as well as to the specific steps of the staining protocol^94,95^. Additionally, there are also non-adherent biofilms, e.g. those at the air-surface interface or as floating “flakes” (e.g., for Alteromonas^45^). While the crystal violet assay can, in principle, be used to measure non-adherent biofilms (e.g. using filtration), it will require adaptation and tuning, especially if applied across diverse bacteria.

## Discussion and Conclusions

Bacterial diversity is truly astounding. Each strain has its own unique “personality” - the shape of its growth curve, the color and shape of the colonies, or the way it swirls, aggregates or adheres to the test tube when shaken. These idiosyncratic observations of the lifestyle of each bacterium may be related to organismal or ecosystem-level traits that are important in specific environments, such as growth rates on different resources or the ability to swim towards and colonize new niches (e.g. hosts or particles). As a crucial step towards understanding the diversity of these traits and their genetic underpinning, we have established a collection of bacteria that can be cultured under similar conditions, and a set of assays that can be applied systematically across them. Applying well-established methods across such diverse culture collection proved to be challenging, as illustrated by the discrepancies between standard methods for counting cells and measuring their motility. Future work will require harmonizing the systematic viewpoint afforded by culture collections and high-throughput assays with a recognition of the unique and nuanced biology of each organism.

Even though genomic information was used to select the organisms for this library in the first place^28^, we did not attempt systematic evaluations of correlations between genetic features of the library strains and their phenotype in the various assays. Many of the phenotypes examined are likely polygenic and would require much higher statistical power to detect meaningful associations. Our primary goal was to establish a framework for characterizing phenotypic diversity rather than to exhaustively map genotype–phenotype links at this stage. Some assays may still benefit from refinement to better capture phenotypic variability across strains (e.g. cell counts), while others, such as motility and biofilm formation, are influenced by conditions that are difficult to standardize across experiments^97^. Moreover, several genes remain unknown or poorly annotated. For instance, when testing the association between hemolytic activity on blood agar and the number of genes encoding putative hemolysins, we found that 5 of 7 strains encoding 3 or more hemolysins displayed complete (beta) hemolysis, as did 5 strains lacking such genes. These observations highlight both the complexity of inferring function from sequence alone and the value of broad phenotypic screens in revealing unanticipated biological capabilities.

The insights gained from working with strain libraries using robust, standardized and reproducible methods will be critical as we aim to understand “which organism does what” in complex and dynamic microbial communities. Standardized collections of bacterial strains and assays provide a powerful foundation for deciphering microbial lifestyles and functions. Although genomic information guides much of today’s biology, interpreting genomes in functional terms remains a major challenge. Integrating high-throughput phenotypic analyses with genomic data, as highlighted in our recent work^98^, will be key to bridging this gap. The broad adoption of standardized libraries and consistent data-sharing practices across research groups can create valuable synergies, especially for non-model organisms that remain poorly characterized. Moreover, as microbial collections expand in diversity and scale, even seemingly simple assays can become difficult to manage and compare. Systematically documenting and analyzing these methodological challenges is itself informative, as it reveals the complexity of microbial behavior and highlights opportunities for improving analytical approaches. Ultimately, increasing the dimensionality and comparability of phenotypic data will not only enhance reproducibility but also enable a deeper and more integrative understanding of bacterial diversity and function.

## Materials and Methods

Library strains were maintained under constant light in Difco Marine Broth 2216 or on Marine Agar at 26°C. Liquid cultures (2 ml in 5 ml snap-cap tubes) were shaken at 60 RPM for routine maintenance. Cryopreservation was performed by flash-freezing cultures in 50% glycerol upon visible turbidity or after 36–48 h; samples were stored in sterile cryo-tubes or 96-well plates at −80°C. For experimentation, frozen strains were revived by inoculation into fresh Marine Broth, grown for 36–48 h, then subcultured prior to use.

DNA was extracted from exponentially growing liquid cultures using the PureLink Genomic DNA Mini Kit. For 16S rRNA gene amplification, the 27F/1492R primers were utilized, followed by Sanger sequencing and BLASTn comparison against the NCBI nr database. Genome resequencing incorporated Illumina and Oxford Nanopore platforms; mutations were identified using the *breseq* pipeline with default settings.

Growth assays were performed in 96-well plates (200 µl per well) at temperatures ranging from 10–30°C. OD600 was measured at specified intervals, with condensation controlled via sterile hood drying. Chemically defined media substituted complex components with amino acids, peptides, sugars, and organic acids at specified concentrations to assess substrate utilization. Absorbance fold-change was evaluated relative to negative controls. Flow cytometry and CFU assays were performed in biological and technical replicates, following established protocols. Hemolysis, antibiotic susceptibility (Kirby–Bauer method), motility, and biofilm formation (crystal violet staining) assays were conducted with standardized conditions and measurements. All data and scripts are publicly available. See supplementary information 1 for detailed materials and methods.

## Abbreviations

rpoB: RNA polymerase beta subunit
rpoC: RNA polymerase beta’ subunit
ssb1: single-stranded DNA binding protein 1
GFCs: Genome Functional Clusters
KEGG: Kyoto Encyclopedia of Genes and Genomes
DMSZ: Deutsche Sammlung von Mikroorganismen und Zellkulturen
ATCC: American Type Culture Collection
MB: Marine Broth
MA: Marine Agar
mM: millimolar
mL: milliliter
GC: guanine-cytosine content
SNPs: single nucleotide polymorphisms
dN/dS: ratio of non-synonymous to synonymous substitutions
OD600: optical density at 600 nm
LB: Luria Bertani Broth (also known as Lysogeny Broth)
C: carbon
CCR: carbon catabolite repression
FCM: flow cytometry
CFU: colony forming unit
qPCR: quantitative polymerase chain reaction
DAPI: -6,’4diamidino-2-phenylindole
Sybr Green: SYBR Safe DNA Gel Stain (brand of nucleic acid stain)
ROS: reactive oxygen species
DNA: deoxyribonucleic acid
RPM: revolutions per minute
PCR: polymerase chain reaction
NCBI: National Center for Biotechnology Information
nr: non-redundant database
µl: microliter
FDR: false discovery rate

## DECLARATIONS

### Ethics approval and consent to participate

Not applicable

### Consent for publication

Not applicable

### Availability of supporting data

Raw sequencing reads are available in the NCBI Sequence Read Archive (SRA) under BioProject PRJNA1367194. The breseq codes have been made available in Zenodo at https://doi.org/10.5281/zenodo.17584611, and all reference genomes are also available at https://doi.org/10.5281/zenodo.17584547. All python scripts used to generate various plots for the figures, along with their data and the statistics used is available on the <u>Sher Lab GitHub</u>.

### Author Contribution

WBVK, LZ, MO, EF, DSe, HPG and DSh conceived the study and selected the strains for the library. WBVK and DSh analyzed the data and wrote the paper. EF performed the hemolysis Assay and WBVK performed all other experiments and assays. MO and EF collected and maintained the strains. FK, MO, and WBVK cultured and processed samples for strain genome resequencing. KH and WBVK performed strain identification by 16S rRNA gene sequencing. HS and MO performed mutation analysis with the help of NG and JV-R. KS and CAMG performed the phylogenetic reconstruction. LZ performed GFC analysis. DSe, HPG and DSh provided funding.

## Acknowledgements

We thank the laboratories of Xuewei Xu, Roberto Kolter, Joseph Christie-Oleza, Francisco Rodriiguez Valera and Mary Ann Moran for some of the strains in the library.

