## Supplementary Information for "Biological insights and methodological challenges learned from working with a diverse heterotrophic marine bacterial library"

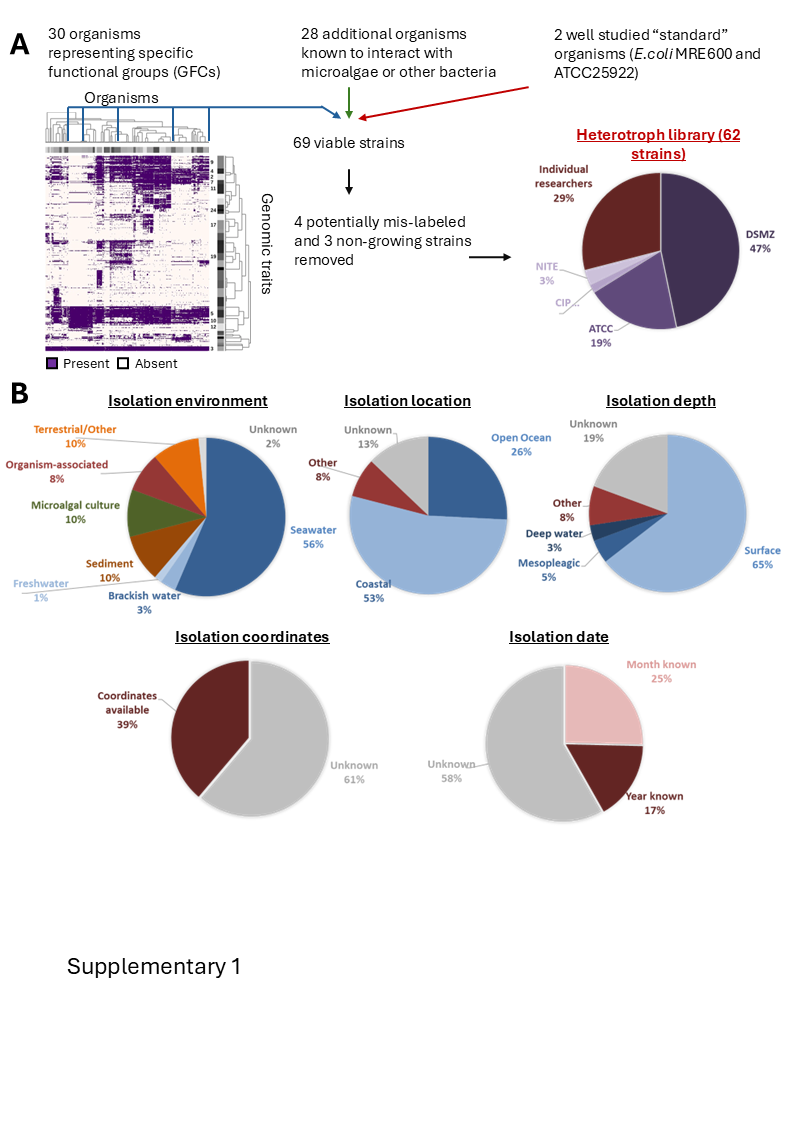

Supplementary Figure S1: General description of the marine heterotroph library. A) Overview of the strain selection process, starting from a map trait across genomes and including also selection of bacteria known to interact with phytoplankton, and well-studied model organisms. The pie chart shows the sources from which the bacteria were obtained. B) Information on the strains in the library. The heatmap in panel 1A is from a previously published manuscript^1^.

-

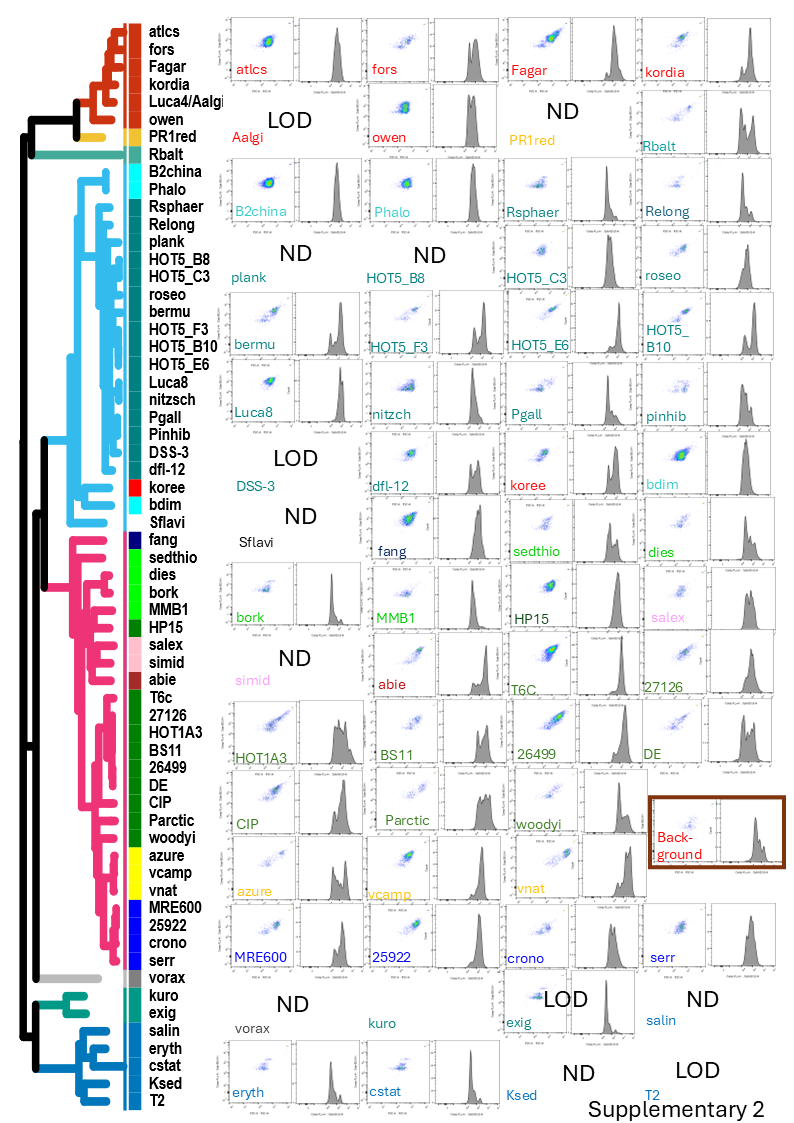

Supplementary Figure S2: Flow cytometry cytograms, and Sybr-staining histograms, of the strains in the library. The strains are organized by their phylogenomic order in the tree. Note the background noise, likely from the Marine Broth.

-

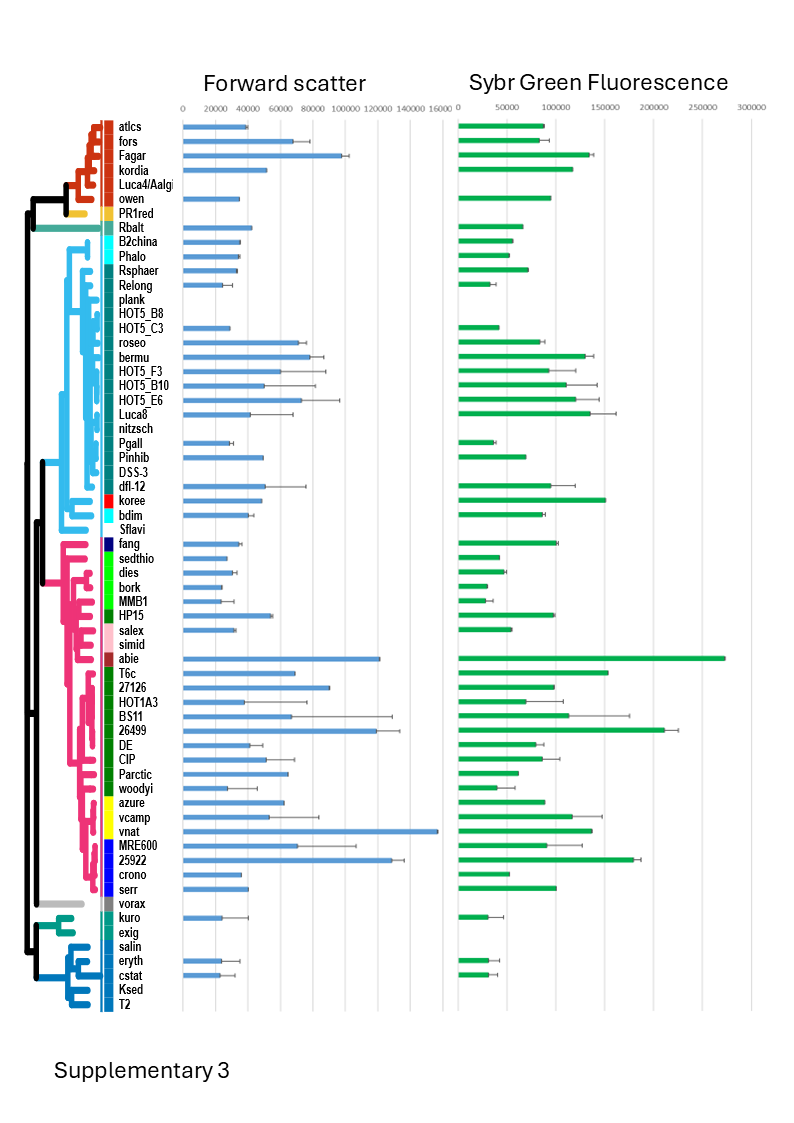

Supplementary Figure S3: Forward scatter and Sybr Green fluorescence across the library strains after 36h growth in Marine Broth. Results shown are mean and standard deviation of the gated cell population.

**DETAILED MATERIALS AND METHODS**

**Library maintenance and preservation.**

The strains in the library were routinely cultured in the light in liquid Difco Marine Broth 2216 or on Marine Agar plates at 26h (the composition of all media used in these experiments is detailed in Supplementary Table 1). Routine maintenance was performed in constantly shaken liquid cultures (2ml liquid in 5ml snap-cap tubes, typically shaken at 60 RPM). Most experiments were performed in 96 well plates, with 200ml media in each well, which were shaken only prior to measurement (every 2-12 hours, as described for each experiment below). Cryo-preservation was performed in 50% glycerol at the earliest point turbidity was seen, or after 36-48 hours. The library was maintained both sterile 2ml cryo-tubes, which were flash-frozen in liquid nitrogen before transfer to a 80ºC freezer, or arrayed in two 96 well plates, which were frozen at -80ºC.

Prior to specific experiments the library strains were revived by scraping the tip of the frozen vials or wells using a sterile pipette tip and inoculating into fresh liquid Marine Broth, either in tubes or 96 well plates. After an initial 36-48 hours, several microliters from these starter cultures were transferred into fresh marine broth for another 36-48 hours growth before starting the experiments.

**16S rRNA gene amplification, genome resequencing and analysis**

DNA was extracted from liquid cultures in MB. For 16S rRNA gene amplification, 5 ml cultures were used, whereas for genome re-sequencing 5 – 15 ml were needed to prepare at least 6 x 10^9 cells; The pellets were gently resuspended in 500 µL of Zymo 1X DNA/RNA Shield preservative (R1100) before sending because some genome re-sequencing attempts were initially unsuccessful, we typically attempted to extract DNA from exponentially growing cultures. DNA was extracted using the PureLink™ Genomic DNA Mini Kit. For 16S amplification, the 27F and 1492R primers^2,3^ were used, followed by Sanger sequencing (Macrogen Europe). The sequences were then compared to the ncbi nr database using BLASTn.

Genome re-sequencing was performed using a combination of Illumina and Oxford Nanopore technologies (Pasmidsaurus; culturing, pelleting and resuspending was done as previously mentioned). Average Nucleotide Identity was assessed between the re-sequenced and reference genomes using skani^4^ and mutations were identified using *breseq*^5^, an open-source pipeline that automates all steps of the variant calling pipeline from mapping reads to a reference genome, to annotation, and is optimized for use on haploid microbes. All default settings were used to run *breseq*, and the consequence of mutations were added into an annotated GenomeDiff format using the gdtools command installed as part of *breseq*.

**Phylogenomic reconstruction and association with previous Genome Functional Clusters**

In order to evaluate the phylogenetic breath of the strains in our collection, we built  phylogenomic trees using 30 universal marker genes belonging to RNA polymerase subunits and Ribosomal proteins through the pipeline MarkerFinder^6^. Protein-predicted files generated with Prodigal V2.6.3^7^ were used as input for MarkerFinder, and further steps consisted in marker gene annotation with hmmsearch (HMMER 3.2.1)^8^ and their alignment with Clustalo 1.2.4^9^ using default parameters. The output concatenated alignment was trimmed with TrimAl using -gt 0.1^10^. Tree reconstruction was performed through IQ-TREE v1.6.9^11^ with the options -bb 1000 (bootstrap replicate generation^12^) and -MFP for substitution model selection under the BIC criterion with ModelFinder^13^. The final topology was explored on iTOL to discard the occurrence of topological biases^14^.

To integrate the strains from the library into the previously described framework of Genome Functional Clusters^1^ (GFCs), we annotated the genomes not included in the previous study based on gene calling by Prodigal v2.6.3^15^ and subsequent annotation using a snakemake workflow (available at <https://github.com/lucaz88/FunTraits>). Module completeness for the thiamine (M00127) and cobalamin (M00122) KEGG pathways was manually validated, and only transmembrane proteins associated with vitamin and siderophore transport were retained. To merge the new annotation with the previous GFCs, pairwise Pearson correlations were computed between binary functional trait profiles, negative correlations were set to zero and only pairwise correlations with a FDR-corrected p-value < 0.05 (chi-square test, df = 1) were retained. Affinity propagation clustering was applied to correlation matrices (apcluster function, q = 0.5) (R package apcluster 1.4.8^5^; Bodenhofer et al., 2011) to infer GFCs de novo. Concordance between new and reference GFCs was assessed by mapping cluster membership. GFC labels were propagated according to three criteria: (i) if a new GFC perfectly matched a reference GFC or differed only by the inclusion of new isolates, the original reference label was retained; (ii) if a new GFC comprised exclusively novel genomes with no correspondence to reference clusters, a new label was assigned (marked with ###); and (iii) if genomes assigned to distinct new GFCs mapped to the same reference GFC, each cluster was assigned a unique label (e.g., NEWb_##, NEWc_##, etc.). Newly assigned clusters arising from cases (ii) and (iii) were reviewed, and those deemed likely to represent technical artefacts rather than biologically meaningful groupings were excluded from downstream analyses.

**Growth at different temperatures**

The starter plates described above were each transferred 1:100 v/v into fresh 96 well plates with media preconditioned to the experimental temperatures (10, 15, 20, 25 and 30ºC). Thus, this experiment measures the immediate response to temperature shifts rather than the response of pre-acclimated strains. Culture turbidity (OD_600 nm_) was then measured every ~2 hours for 24 hours. Because of the differences between the temperature in the incubator and the room where the measurements were performed, condensation was observed on some plates. In such cases the plate lid was removed and dried for a few minutes inside a sterile biological hood before reading the plates. After manual cleaning of the data from clear outliers, growth rate was calculated using the formula Slope (m) = y2-y1/x2-x1, where y is the OD_600_ value at time (x). The maximal slope was calculated and taken for further visualization.

**Growth in specific defined media**

Marine Broth is a complex media containing yeast extract and peptone, which are not chemically defined. We previously re-factored marine broth to replace these components with chemically defined molecules (^16^ , all media compositions are described in the Supplementary Excel File). We replaced Fe-citrate in the salts base of the refactored MB with Ferric Chloride to remove this potential organic carbon source. Alpa-Casein was used as a source of protein, peptides (trypsinized) and free amino acids, at a concentration of 3g/l. For individual amino acids this results in concentrations in the range of 16uM (L-glutamate) and 1.3 uM (L-tryptophan). Because tryptophan is degraded during the acid hydrolysis step used to make amino acids, it was added separately (0.5 g/l). Media which included also carbohydrates had a mixture of sugars (0.166 g/l of Maltose, Trehalose and Sucrose) and organic acids (0.284 g/l each of Lactate, Pyruvate, Acetate, Citrate, Succinate and Glycolate) as tabulated. While all media was adjusted to a neutral pH, for the whole protein media, pH was not adjusted to avoid curdling.

The library was inoculated into 96 well plates containing the different media as described above for the temperature experiment, and absorbance at 600nm was read approximately every 12 hours for a total of 60 hours. The absorbance of blank wells was subtracted, and the fold-change in absorbance was calculated compared to that of the negative control for each strain (i.e. grown on marine broth base without any carbon sources). Strains where no growth was observed in the positive control (refactored marine broth) were removed from the analysis.

**Comparison of flow cytometry and colony forming units**

The comparison of FCM and CFU was performed three separate times (biological replicates), each with technical replicates (wells of each strain). The library was grown in 96 well plates with 200ul MB without shaking as described above for 36 hours. For flow cytometry, the library was diluted 1:10,000 in MB, fixed with 1ul of 25% glutaraldehyde per well, and maintained in -80°C until analysis. The cells were thawed, 20ul of which was mixed with 5ul of Sybr and 5 ul of beads, both from a diluted stock (20%v/v). Analysis was performed using FlowJo version 10.8.0. For CFU counts, the library was first serially diluted 1:10 in MB. 10ml each of the 1:10^6^-1:10^8^ dilutions were then pipetted onto the surface of a Marine Agar plate using a multi-channel pipette. Once the droplets were absorbed into the agar surface, the plates were incubated at 22°C for 48 hours or longer until colonies were clearly formed.

**Hemolysis, antibiotic sensitivity, motility and biofilm formation**

Liquid cultures in MB were streaked on commercial Sheep’s Blood Agar plates (Hardy), and the hemolytic phenotype recorded after at earliest observed growth For the Kirby Bauer method for antibiotics sensitivity, 2.5% Marine Agar plates were swabbed thoroughly with a sterilized cotton swab soaked in bacterial culture. Disks containing clinical concentrations of the seven antibiotics (Oxoid™) were then carefully placed on the agar with sterile forceps. Ampicillin (CT0004B -25μg), Nitrofurantoin (CT0034B- 100μg), Tetracycline (CT0053B -10μg), Ciprofloxacin (CT0425B -5µg), Chloramphenicol (CT0012B - 10μg), Kanamycin (CT0025B - 5µg), and Streptomycin (CT0047B -10μg) were chosen. The size of the halo surrounding each disk was recorded after growth at 26°C from 48 hours to 96 hours depending on bacterial growth rate.

Swarming motility was assessed under aerobic conditions by spotting semi-solid marine agar plates (0.5% agar) with 10ml of a liquid culture in the middle of the plate. For motility under anaerobic conditions, a semi-solid agar slant tube was prepared, and the slanted surface was first streaked using a loop of the cultured broth. The streak was then stabbed to inoculate the bacteria into the agar slant^17,18^. Motility in both assays was recorded from 48 to 96 hours after incubation at 26°C.

Biofilm formation was assessed by growing the library in 96 well (100ul) plates for 76 hours at 26°C. Additionally, 1.25X the volume of crystal violet (0.1% w/v working stock) as opposed to media was added and the plates were incubated for another hour. The plates were turned around and shaken until all un adhered mass comes off and rinsed with DDW by shaking without pipetting. The remaining crystal violet was dissolved in acetate. Transfer dissolved acetate (20ul) to fresh plate, diluting it in DDW (180ul) by an order of magnitude. OD550 was measured at room temperature.

**MOTILITY LITERATURE**

| **Strain** | **Name** | **Literature Motility** | **Reference Number** |
| --- | --- | --- | --- |
| cstat | Corynebacterium stationis | N | ^19^ |
| Ksed | Kytococcus sedentarius | N | ^20^ |
| eryth | Rhodococcus erythropolis | N | ^21^ |
| salin | Salinispora tropica | N | ^22^ |
| T2 | Aeromicrobium marinum | N | ^23^ |
| Sflavi | Sphingorhabdus flavimaris | N | ^24^ |
| B2china | Pelagibacterium halotolerans | M | ^25^ |
| Phalo | Pelagibacterium halotolerans | M |  |
| dfl-12 | Dinoroseobacter shibae | M | ^26^ |
| nitzsch | Sulfitobacter pseudonitzschiae | N | ^27^ |
| plank | Planktomarina temperata | M | ^28^ |
| pgall | Phaeobacter gallaeciensis | M | ^29^ |
| pinhib | Phaeobacter inhibens | M | ^30^ |
| koree | Ponticaulis koreensis | M | ^31^ |
| DSS-3 | Ruegeria pomeroyi | M | ^32^ |
| roseo | Roseovarius mucosus | N | ^26^ |
| Rsphaer | Rhodobacter sphaeroides | M | ^33^ |
| bdim | Brevundimonas diminuta | M | ^34^ |
| Relong | Roseibacterium elongatum | N | ^35^ |
| bermu | Pelagibaca bermudensis | N | ^36,37^ |
| exig | Exiguobacterium oxidotolerans | M | ^38^ |
| kuro | Halobacillus kuroshimensis | N | ^39^ |
| PR1red | Algoriphagus machipongonensis | N | ^40^ |
| owen | Owenweeksia hongkongensis | M | ^41^ |
| atlcs | Croceibacter atlanticus | N | ^42^ |
| Fagar | Formosa agariphila | M | ^43^ |
| fors | Gramella forsetii | M | ^44^ |
| Luca4/Aalgi | Arenibacter algicola | N | ^45^ |
| kordia | Kordia algicida | N | ^46^ |
| bork | Alcanivorax borkumensis | N | ^47^ |
| dies | Alcanivorax dieselolei | M | ^48^ |
| 26499 | Alteromonas mediterranea | M | ^49^ |
| 27126 | Alteromonas macleodii | M |  |
| BS11 | Alteromonas macleodii | M |  |
| DE | Alteromonas mediterranea | M |  |
| HOT1A3 | Alteromonas macleodii | M |  |
| T6c | Pseudoalteromonas atlantica | M | ^50,51^ |
| abie | Pseudomonas abietaniphila | M | ^52^ |
| 25922 | Escherichia coli | M | ^53^ |
| crono | Cronobacter universalis | N | ^54^ |
| MRE600 | Escherichia coli | N | ^55^ |
| serr | Serratia sp. | M | ^56^ |
| fang | Fangia hongkongensis | N | ^57^ |
| salex | Haliea salexigens | M | ^58^ |
| HP15 | Marinobacter adhaerens | M | ^59^ |
| MMB1 | Marinomonas mediterranea | M | ^60^ |
| CIP | Pseudoalteromonas haloplanktis | M | ^50^ |
| sedthio | Sedimenticola thiotaurini | M | ^61^ |
| HOT5_B8 | HOT5_B8 | M | - |
| HOT5_E6 | HOT5_E6 | M | - |
| woodyi | Shewanella woodyi | M | ^62^ |
| simid | Simiduia agarivorans | N | ^63^ |
| azure | Vibrio azureus | M | ^64^ |
| vcamp | Vibrio campbelli | M |  |
| vnat | Vibrio natriegens | M |  |
| Parctic | Psychromonas arctica | M | ^65^ |
| Luca8 | Luca8 | M | ^27^ |
| vorax | Pseudobacteriovorax antillogorgiicola | M | ^66^ |
| HOT5_B10 | HOT5_B10 | M | - |
| HOT5_C3 | HOT5_C3 | M | - |
| HOT5_F3 | HOT5_F3 | M | - |
| Rbalt | Rhodopirellula baltica | M | ^67^ |

Abbreviations: M: Motile, N: Non-Motile.

Empty cells are unpublished isolates still under characterization that have been marked as Motile based on other strains that are phylogenetically related.
